# Gradient scheme optimization for PRESS-localized edited MRS using weighted pathway suppression

**DOI:** 10.1101/2025.07.14.664705

**Authors:** Gizeaddis L. Simegn, Zahra Shams, Saipavitra Murali-Manohar, Dunja Simicic, Abdelrahman Gad, Yulu Song, Vivek Yedavalli, Christopher W. Davies-Jenkins, Aaron T. Gudmundson, Helge J. Zöllner, Georg Oeltzschner, Richard A.E. Edden

## Abstract

This study aimed to design and implement an optimized gradient scheme for PRESS-localized edited magnetic resonance spectroscopy (MRS) to enhance suppression of out-of-voxel (OOV) artifacts. These artifacts, which originate from insufficient crushing of unwanted coherence transfer pathways (CTPs), are particularly challenging in editing schemes for metabolites like gamma-aminobutyric acid and glutathione. To address this, a volume-based likelihood model was developed to guide gradient scheme optimization, prioritizing suppression of CTPs based on likelihood.

A volume-based likelihood model for CTP weighting was integrated into a Dephasing optimization through coherence order pathway selection (DOTCOPS) gradient optimization. Using a genetic algorithm with a new dual-penalty cost function, gradient schemes were optimized to maximize pathway-specific suppression. Hardware and sequence constraints, maximum gradient amplitudes and delay durations respectively, informed the optimization. Validation of the optimized scheme was performed with simulations by calculating the k-space crushing efficiency analytically with k-space trajectory. and *in vivo* using an edited MRS sequence in three brain regions (posterior cingulate cortex PCC, thalamus, and medial prefrontal cortex (mPFC)), with particular focus on OOV artifact reduction and spectral quality improvements. A three-way Analysis of Variance was used to assess the significance level of OOV artifact reduction.

The optimized gradient scheme demonstrated improved k-space crushing efficiency (by an average of 197%). OOV artifacts were reduced in all brain regions, particularly in highly OOV-susceptible regions (thalamus and mPFC). Improvements were most notable around 4.3 ppm with significant OOV artifact amplitude reductions (p < 0.001).

By using a volume-based likelihood model for CTP prioritization, the optimized DOTCOPS scheme ensures robust and region-agnostic performance in reducing OOV artifacts.

## 1. Introduction

Proton magnetic resonance spectroscopy (^1^H MRS) is a non-invasive imaging modality that measures the *in vivo* concentration of metabolites in brain and body tissues, such as N-acetyl aspartate (NAA), choline-containing compounds (Cho), glutamate, glutamine, lactate, and others, which are potential biomarkers for conditions ranging from brain tumors to psychiatric disorders (1–6). MRS experiments are implemented as pulse sequences of radiofrequency (RF) and magnetic field gradient pulses, designed to control the evolution of the intended signals from the localized region of interest and suppress other potential nuisance signals. One useful tool for keeping track of the evolution of a given coherence through the pulse sequence is a coherence transfer pathway (CTP) diagram, which plots the coherence order at each stage of the sequence (7). Unwanted signals generally arise from different CTPs from the intended signals and degrade spectral quality and quantification accuracy (8,9). Among the most notorious of those nuisance signals are out-of-voxel (OOV) signals, spurious echoes from water signals outside the voxel refocused by local field gradients (10).

OOV signals represent spurious echoes from water protons outside the voxel of interest that are inadvertently refocused by local magnetic field gradients, particularly near tissue boundaries. In our recent study (11), the dominant OOV contribution was traced to a specific coherence transfer pathway (Δp = [0, 0, 0, 0, –1]) excited by an editing pulse and not subsequently refocused, rendering it resistant to conventional phase cycling. Spectrally, they manifest as broad features mainly between 3.9–4.5 ppm. In general, they are more likely to appear between 3.9 and 4.5 ppm because that is close to the shimmed frequency of the water resonance (OOV signals the other side of water often being present, but not visualized), but they can appear at any frequency. In the case of the ‘edit-excite’ pathway that drives many of the signals seen in these data, the editing pulse lobes at 4.58 ppm and 4.18 ppm are at fault. It is important to note that while this pathway was a major contributor in that specific context, the prevalence of different artifact-generating CTPs can vary; our current optimization framework is therefore designed to be general and accounts for the likelihood of multiple unwanted pathways.

Low-concentration metabolites like γ-aminobutyric acid (GABA) are often measured with J-difference-edited MRS experiments (12) such as MEGA-PRESS (13) and Hadamard Editing Resolves Chemicals Using Linear-combination Estimation of Spectra (HERCULES) (14). CTP selection is particularly critical in edited MRS both because measurements of weak signals are less tolerant of small artifacts, and because edited MRS experiments have more RF pulses and sub-experiments than their unedited counterparts, increasing the number of CTPs to be controlled exponentially. For instance, in a standard five-pulse sequence such as MEGA-PRESS, considering only coherence orders of ±1 and 0, there are 3 = 81 potentially detectable CTPs. This number arises because each of the first four RF pulses can yield three coherence orders (p = +1, 0, or –1), and only pathways that conclude with a detectable p = –1 coherence after the fifth pulse are observed. Among these 81 possible pathways, only one corresponds to the desired edited metabolite signal, while the remaining 80 represent potential sources of artifacts.

Pulsed field gradients (“crusher gradients”) are the primary mechanism for suppressing unwanted CTPs (15–17). These pulses, applied during evolution delays between RF pulses, impart phase shifts to coherences that depend on location and coherence order. An effective crusher gradient scheme results in the intended CTP being refocused, while unwanted CTPs accumulate destructive phase dispersion across the volume, effectively eliminating their signal contribution (18). Conventional gradient scheme designs often rely on heuristic strategies—iterative adjustments of gradient timing, duration, and amplitude until artifact-free spectra are empirically achieved. While this trial-and-error approach is feasible for basic localization sequences (e.g., PRESS or STEAM (19)), it becomes inadequate for advanced edited sequences due to the sheer number of pulses and gradients involved.

More systematic strategies, such as the Dephasing Optimization Through Coherence Order Pathway Selection (DOTCOPS) scheme (20,21), have formalized the design of crusher gradient schemes by using an algorithmic approach. DOTCOPS represented a significant advance over the traditional approach for MRS gradient scheme design for dephasing unwanted CTPs. The DOTCOPS optimization function considers all unwanted CTPs, using the ‘k-space dephasing distance’ as a proxy for pathway suppression, and seeking to maximize suppression of all pathways on average, and the least-suppressed pathway specifically. Although the option of differential weighting of pathways was suggested, in the absence of a theoretical basis to be more concerned with one CTP than another, no weightings were applied. In reality, not all unwanted CTPs pose the same risk to spectral quality, because some CTPs are more likely to be excited than others for a given pulse sequence configuration, and a model from which to calculate such weightings is required.

In this work, we introduce a volume-based model for CTP likelihood, and incorporate it into a weighted DOTCOPS gradient scheme optimization for PRESS-localized edited MRS. The proposed method introduces a pathway-specific suppression strategy, where CTPs are prioritized based on their likelihood of contributing to spectral contamination. This theory-guided weighting allows for a more intelligent allocation of gradient amplitudes, focusing suppression efforts on pathways that pose a greater risk to spectral quality and away from pathways that are improbable. The optimization is further guided by a new cost function that balances pathway suppression, emphasizing prioritized CTPs in both the minimum and average regularized suppression terms across all CTPs for globally consistent performance.

## 2. Methods

This work presents a gradient scheme specifically designed and optimized for PRESS-localized, MEGA-edited spectroscopy. The optimization incorporated a volume-based model for CTP likelihood into a weighted DOTCOPS framework.

### 2.1. Coherence Transfer Pathway (CTP) likelihood model

All possible CTPs were systematically computed based on the sequence design, which includes five RF pulses. For each pulse, coherence orders were tracked, leading to 81 detectable pathways, representing all potential signal trajectories through the sequence. Each pathway’s likelihood of contributing to the final acquired signal was estimated, and these values were used as weights during the gradient optimization. Pathway likelihoods were determined using a theoretical weighting scheme based on the expected contribution of each CTP to unwanted spectral contamination.

The likelihoods of each CTP were estimated by classifying the transition induced by each RF pulse based on the change in coherence order (Δ*p*). Transitions with Δ*p* = ±2 correspond to the action of 180° refocusing pulses, Δ*p* = ±1 correspond to 90°-like pulses, and Δ*p* = 0 lying outside the band of action, based on the principles of spin dynamics (22,23). These pulse behaviors were then mapped onto specific spatial and spectral regions for slice- and frequency-selective pulses, respectively.

To model the likelihood of different CTPs, we classified tissue regions based on their experience of each RF pulse, simplifying their spatial selectivity profiles into discrete zones. For the slice-selective excitation pulse, regions were treated as either “within-slice” or “out-of-slice”, corresponding to experienced flip angle of 90° or 0°, which we associate with CTPs of Δ*p* = ±1 and Δ*p* = 0, respectively. The excitation pulse is modeled as slice-selective pulse with a realistic profile, simplified into a binary model. For slice-selective refocusing pulses, regions are classified as “within-slice,” “slice-edge,” or “out-of-slice”, corresponding to CTPs of Δ*p* = ±2, ±1, and 0, respectively. For editing pulses, analogous categories were defined, “within-edit,” “edit-transition,” and “edit-off,” representing on-resonance, band-edge, and off-resonance effects. Relative probabilities were then assigned to reflect the extent of each region, reflecting its probability of contributing signal (see Figure 1). The values were based on the spatial selectivity profiles of the RF pulses (typical characteristics of RF pulses) and the typical scenario of a voxel that is substantially smaller than the brain. Specifically, “within-slice” and “within-edit” transitions were assigned a baseline relative probability of 1. For slice-edge transitions, a lower probability of 0.2 was assigned reflecting the smaller extent of the transition band relative to the passband. In contrast, the large out-of-slice regions, which do not experience the RF pulse but may still undergo coherence evolution and refocusing (24), were heavily weighted with a probability of 10. For editing pulses, both within-edit and edit-transition spins were assigned equal probabilities of 1 due to the smoother transition profile of editing pulses which produce less rectangular inversion bands. Edit-off regions, being outside the editing band, were assigned a probability of 5, reflecting the greater extent of the chemical shift range relative to the editing pass band. We have chosen to represent these probabilities with estimates so they can be generally applied, rather than making accurate calculations of for one particular slice profile and voxel/head size.

**Figure 1:**
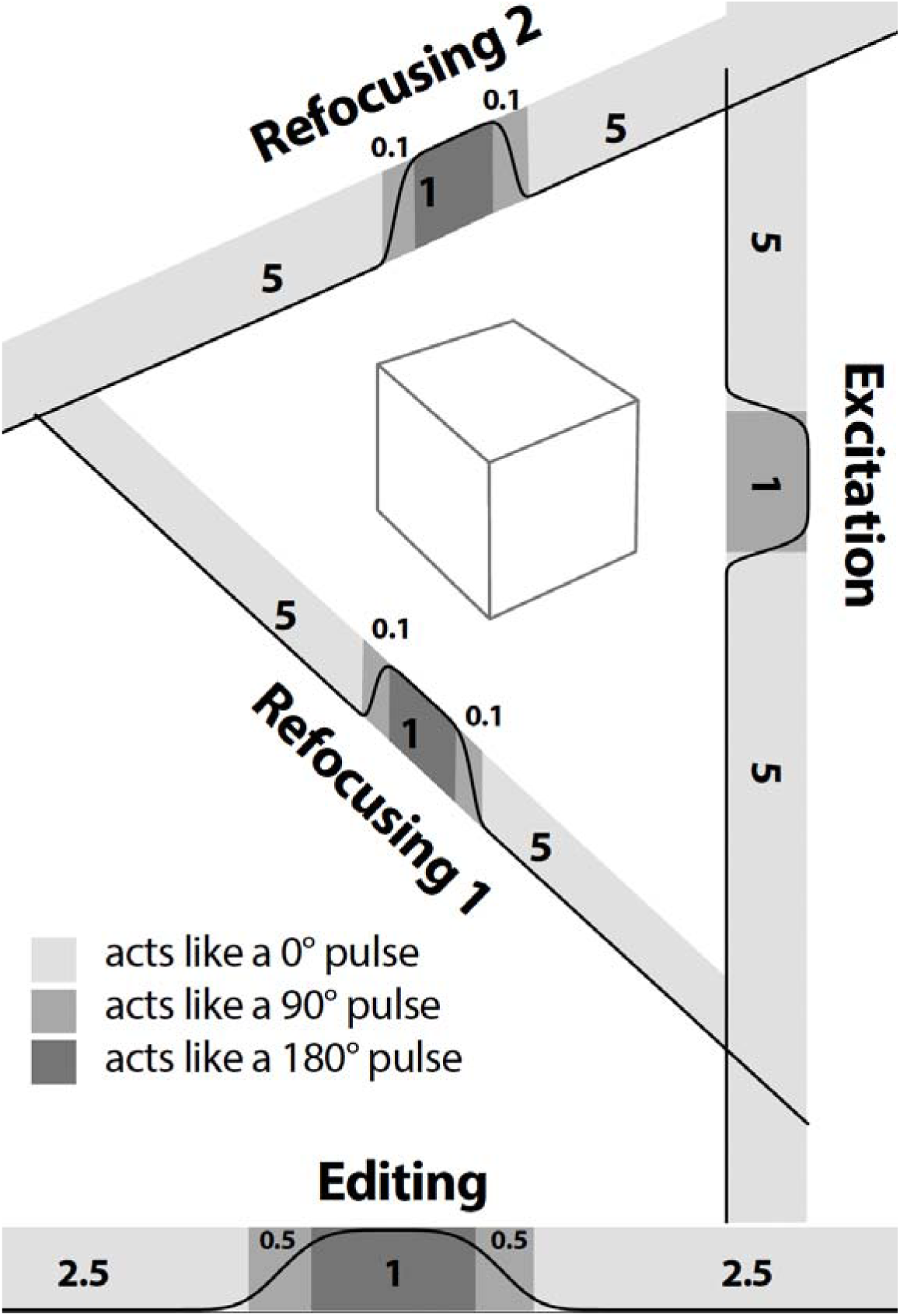
Estimation of CTP likelihoods based on the spatial and spectral profiles of RF pulse-induced transitions. For slice-selective pulses, transitions occurring within the excited region (“within-slice”) are modeled as refocusing-like (Δ*p* = ±2) and assigned a relative probability of 1. Regions within the excitation pulse passband and transitions at the edge of refocusing and editing pulses experience 90°-like pulses (associated with CTPS of Δ*p* = ±1). The total transition-band probability is 0.2 for the refocusing pulse and 1 for the editing pulse. Out-of-band ‘transitions’ of Δ*p* = 0 are given a higher likelihood of 10 for slice-selective pulses and 5 for editing pulses, reflecting the relative extent of each region. Note that transition-band and out-of-band regions occur on either side of the passband of each pulse, so the probabilities on each side are half the total likelihood values, respectively. Pulse profiles are for illustration purposes and are not accurate simulations.

Please note that, the probabilities assigned are generalized estimates applied uniformly across each pulse types in the sequence to ensure a single optimization that applies to all sub-experiments.

The relative likelihood L(*i*) of a CTP *i,* consisting of the transitions *T_i,n_* (i.e., Δ*p* = 0, ±1, ±2 occurring at pulse *n* in pathway *i*) in an N-pulse sequence then becomes the product of these transition probabilities as shown in Equation 1:

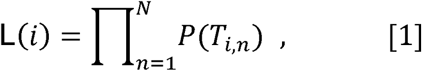

in which, *P*(*T_i,n_*), is the probability weight assigned to that transition, based on spatial or spectral behavior (“within-slice”, “edge”, or “out-of-slice”), as summarized in Figure 1.

For the case of a 5-pulse sequence with 81 possible CTPs, corresponding to 405 total pathway transitions, almost half of all transitions are Δp = ±1, which is the lowest-likelihood action of every pulse. If all pathways are treated equally in gradient scheme optimization (as in previous work (20,21)), the large number of these low-likelihood CTPs can misdirect the focus of the optimizer. While DOTCOPS was always conceived with the option of weighting CTPs differently, this ‘volume-based likelihood’ model provides a conceptual basis for the weights, allowing the optimization to prioritize the suppression of pathways that are more likely to generate out-of-voxel artifacts.

### 2.2. Gradient scheme optimization

A gradient scheme was designed to increase suppression of unwanted CTPs in MEGA-edited PRESS-localized experiments (including HERCULES (14)), combining theory-driven weights, hardware- and sequence-informed constraints, and global optimization. The gradient durations were first precalculated to their maximum allowable values, determined by the available time in each of the five sequence delays between RF pulses. The genetic algorithm was then tasked with optimizing only the gradient amplitudes for each axis within this fixed-duration framework. The optimization aimed to adjust the gradient amplitudes across three axes (*x, y, z*) within the five sequence delays. Gradient durations were constrained by the maximum available sequence delay between each pair of RF pulses: between the excitation pulse and first refocusing pulse; first refocusing pulse and first editing pulse; first editing pulse and second refocusing pulse; and between the second refocusing pulse and second editing pulse.

A genetic algorithm (GA) (*ga* function in MATLAB’s Global Optimization Toolbox) was used for the optimization process due to its robustness in exploring high-dimensional and non-linear search spaces. The weighting *w_i_* assigned to the *i*^th^ pathway were calculated from *L_i_* according to:

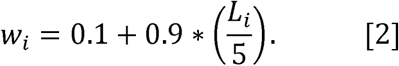

The optimization objective was designed to balance two goals: ensuring sufficient gradient dephasing for the least-suppressed pathways and maintaining overall suppression efficiency as shown in Equation 3:

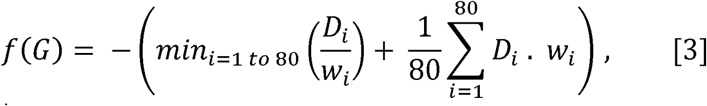

where, *w_i_* is the *i*^th^ pathway normalized weighting and *D_i_* is the L2 norm of net crusher moment for the *i*^th^ pathway expressed as:

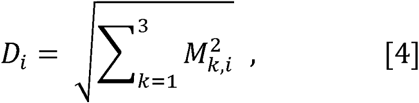

where *M* = *G* × *P* ∊ R^3×80^ is the effective crusher moment for each pathway in each direction *k*, calculated as the matrix product of *G* ∊ R^3×5^, the gradient scheme, and *P* ∊ R^3×80^: the matrix of CTPs). To preserve the intended CTP, equality constraints were applied to balance the gradient moments. Specifically, the sum of the second and third gradient areas in each direction was constrained to equal the combined areas of the first, fourth and fifth gradient areas, in line with refocusing the desired pathway of p = (−1, +1, +1, −1, −1).

While most gradient durations were constrained to fill their respective delays, a 1 ms delay was reserved after the final gradients to mitigate the effects of eddy currents and hardware-induced phase distortions on the acquired signal.

### 2.3. Simulation of K-space crushing efficiency

The method applied to simulate k-space crushing efficiency is an analytical k-space trajectory calculation. This technique models the effect of the crusher gradients by calculating the net k-space displacement (k) imposed on a spin for every possible coherence transfer pathway (CTP). This displacement is the vector sum of the products of each gradient moment (*G_i_*) and the change in coherence order (Δ*_pi_* at each of the five RF pulses in the sequence:

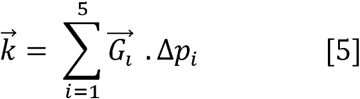

The magnitude of this k-vector (*k_rms_*) is then computed for each pathway. A larger *k_rms_* value indicates a pathway will be more effectively crushed due to significant phase dispersion across a space, while a value near zero indicates the signal will be detectable.

### 2.4. Gradient-scheme-induced diffusion weighting

The incidental diffusion weighting due to the elongated duration of the new crusher schemes is estimated by calculating the *b*-value and signal loss according to reference (25), as in Equation 6:

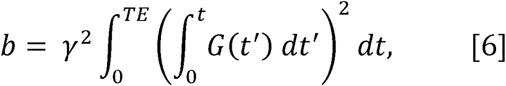

where *γ* is the gyromagnetic ratio (267.522×10 rad/T/s), *G*(*t*) is time-dependent gradient waveform (in *T/m*) and *TE* is the echo time (in *s*). The total b-value, which includes the diffusion due to the interaction of gradient pairs, can be calculated from the self-term (*b_self,i_*: contribution of diffusion weighting from a single isolated gradient pulse) and cross-term contributions (*b_cross,i,j_*: due to the overlapping gradients *i* and *j* on different axes (e.g., *x* and *y*) in different axes):

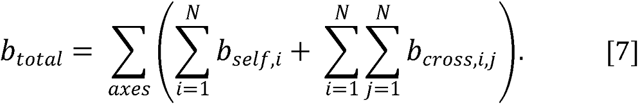

Then, the signal attenuation due to diffusion can be calculated from *b_total_* given the average diffusion coefficient *D_av_* of metabolites or tissue water:

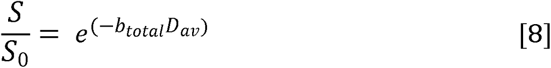

### 2.5. MRS data acquisition

The performance of the optimized gradient scheme was assessed by *in vivo* MRS experiments on ten healthy volunteers (4 females, 6 males, age 26.3±3.4 years). All volunteers provided written informed consent to participate. The Johns Hopkins University Institutional Review Boards approved all study procedures. All scans were conducted on a 3T Philips dStream Ingenia Elition MRI scanner (Philips Healthcare, Best, The Netherlands) using a 32-channel head coil. For MRS voxel placement, a T_1_-weighted structural MPRAGE scan was acquired with the following parameters: TR/TE = 8.0 ms/3.7 ms; flip angle 8°; slice thickness 1.0 mm; 150 slices; isotropic voxel size of 1 mm³, accelerated by compressed sensing with a total acquisition time of 2 minutes and 46 seconds. Following structural imaging, data were acquired using the HERCULES (14) protocol from 30 x 30 x 30 mm^3^ voxels centered on posterior cingulate cortex (PCC), followed by 23 x 30 x 23 mm^3^ voxels centered on thalamus, and 23 x 23 x 23 mm^3^ voxels centered on medial prefrontal cortex (mPFC) (AP × RL × FH) (as shown in Figure 2). Scan parameters for HERCULES were: TR/TE = 2000 ms /80 ms; 192 transients sampled at 2000 Hz with 2048 points; and VAPOR (26) water suppression. For each subject, data were acquired twice — once using the “two-last, increased-area”(11)) and once using the newly optimized proposed (“delay-filling optimized”) gradient schemes. The ‘two-last increased-areas’ gradient scheme is an evolution of our previously developed legacy shared-crusher scheme for HERMES/HERCULES sequences. This modification specifically targets areas of the final crusher gradient pairs along two axes. This results in a larger applied gradient moment (area), significantly enhancing the dephasing of OOV signals. By applying this stronger dephasing power along two axes immediately before signal acquisition, the scheme provides more robust suppression of these specific artifacts compared to its predecessor. Identical voxel positioning and sequence parameters were maintained for both acquisitions to ensure a fair comparison.

**Figure 2:**
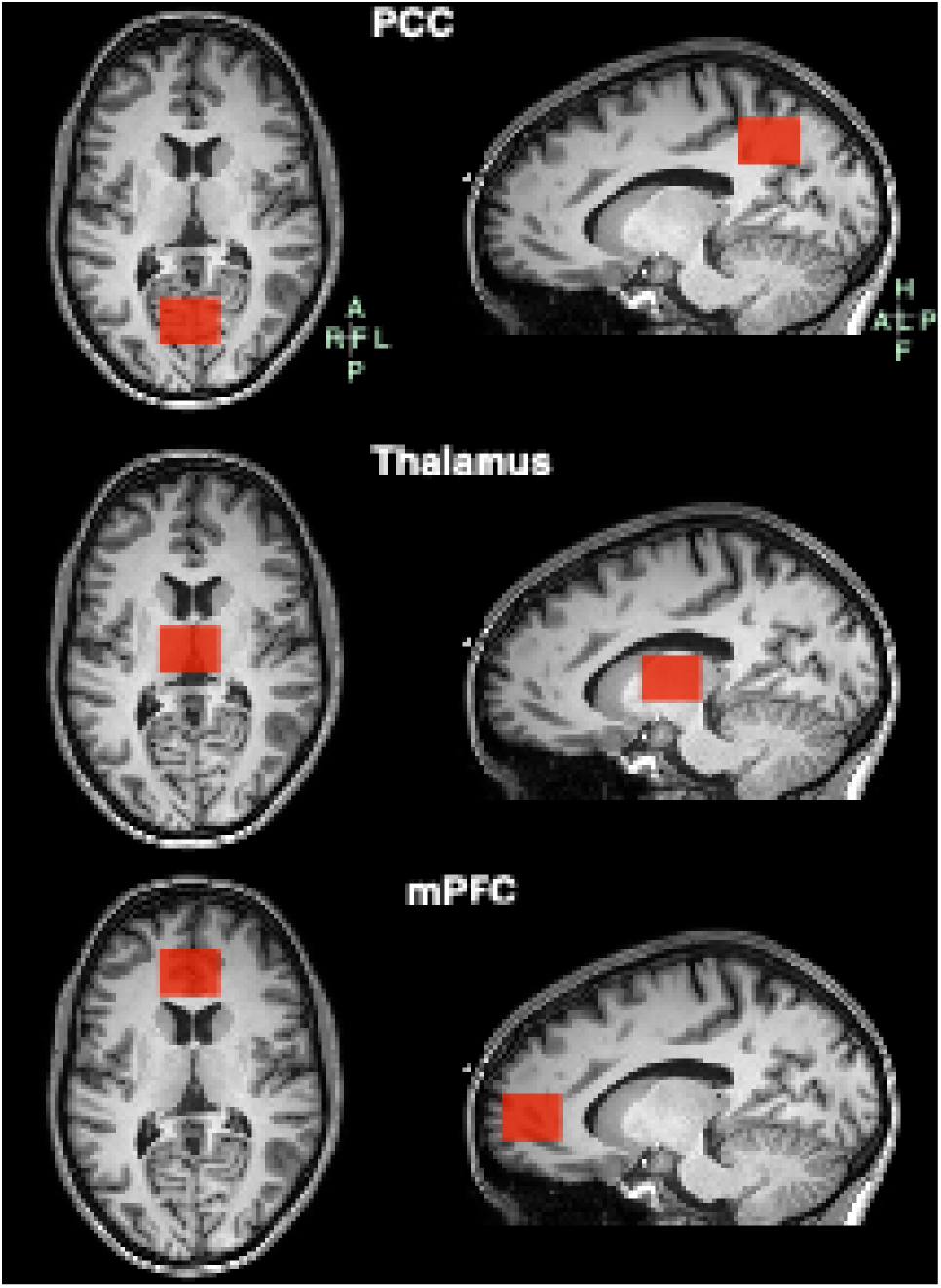
MRS Voxel placement. In vivo edited spectra were acquired from a (30 × 30 × 30) mm^3^ posterior cingulate cortex (PCC), a (23 × 30 × 23) mm^3^ thalamus and a (23 × 23 × 23) mm^3^ medial prefrontal (mPFC) voxels.

### 2.6. MRS data processing and analysis

The edited MRS data were analyzed within MATLAB R2023b using the open-source MRS analysis toolbox Osprey v2.9 (27), following consensus-recommended preprocessing and linear-combination modeling. Frequency and phase correction of FIDs was achieved via robust spectral registration (28). A Hankel singular value decomposition (HSVD) filter was applied to suppress residual water. Linear-combination modeling (LCM) was performed on each of the relevant Hadamard spectrum combinations (i.e., SUM; the sum of all sub-spectra acquired across all sub-experiments, GABA-edited, and GSH-edited difference spectra) from 0.2 to 4.5 ppm, using 17 metabolite basis functions (including Ascorbate (Asc), Aspartate (Asp), Creatine (Cr), Gamma-Aminobutyric Acid (GABA), Glycerophosphocholine (GPC), Glutathione (GSH), Glutamine (Gln), Glutamate (Glu), Myo-Inositol (mI), Lactate (Lac), N-Acetylaspartate (NAA), N-Acetylaspartylglutamate (NAAG), Phosphocholine (PCh), Phosphocreatine (PCr), Phosphoethanolamine (PE), and Scyllo-Inositol (sI), Taurine (Tau)) and 8 macromolecule (MM) and lipid components parameterized as Gaussians for the sum spectrum. The metabolite basis sets were simulated on MRSCloud (29) using a density-matrix approach, which uses use ideal excitation pulses as an approximation to model the evolution of predefined metabolite spin systems (incorporating chemical shifts and J-couplings) and generate accurate, sequence-specific spectral signatures. For GSH-edited spectra, MM peaks at 1.2 and 1.4 ppm were modeled, while GABA-edited spectra included a composite MM basis function with peaks at 3 and 0.93 ppm (30) with a 3 to 2 amplitude ratio. Soft amplitude constraints were applied for MM, lipids, and NAAG/NAA ratios, based on values described in the LCModel manual (31).

### 2.7. OOV artifact quantification

To compare the new and old gradient schemes in reducing artifacts, the relative amplitude of OOV signals for each Hadamard combination spectrum, calculated by Osprey as a quality metric (QM) and quantified using Equation 9, was used (9).

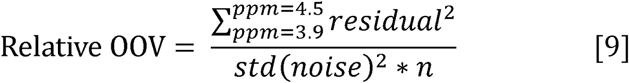

where *residual* refers to the LCM residual between 3.9 to 4.5 ppm (the region of the spectrum with most prominent OOV signals), *std* is the standard deviation, *noise* refers to the real part of noise of the spectrum between −2 and 0 ppm, and *n* is the number of points in the model range. Notably, while this region may include contributions from the broad water suppression tail, its consistent impact across both gradient schemes ensures it does not bias the comparative analysis.

### 2.8. Statistical Analysis

To evaluate whether residuals from OOV echoes differed between the proposed “delay-filling optimized” and the old “two-last, increased-areas” gradient schemes, a multi-way (three-way) analysis of variance (ANOVA) was used. The analysis examined the effects of three factors: ‘Subject’, ‘Voxel location’, and ‘Gradient scheme’ on the mean relative OOV values derived from the SUM, GABA-edited, and GSH-edited spectra. Differences in mean relative OOV values across factor levels were assessed using F-statistics and associated p-values (Prob > F), with statistical significance defined at the 0.05 level.

## 3. Results

### 3.1. Optimized gradient scheme and total crushing moment across CTPs

The proposed “delay-filling optimized” and the “two-last, increased-area” gradient schemes are compared in Figure 3. The genetic algorithm maximized the gradient areas across all three axes, except for the second gradient in each, while reversing the polarity of the first two gradients in the second axis (*G*_y_).

**Figure 3:**
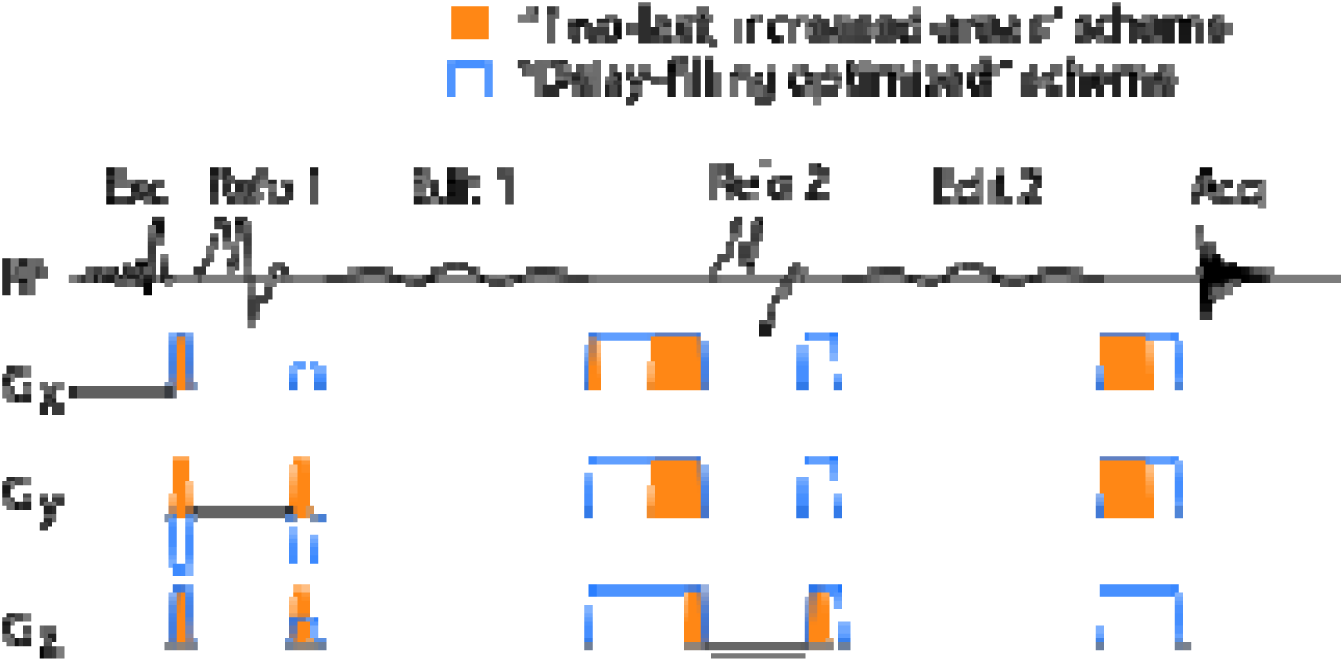
“Delay-filling optimized” (blue) and “Two-last, increased-area” (orange) gradient schemes

The total crushing moment (k-space crushing distance) of each CTP for the “delay-filling optimized” and the “two-last, increased-area” (11), are compared in Figure 4. Each column in the plot represents one of the 81 potential pathways and is aligned with the relevant column of the CTP map, in which where each row indicates the coherence order after a given RF pulse (*RF*_Exc_, *RF*_Refo1_, *RF*_Edit1_, *RF*_Refo2_, *RF*_Edit2_). By convention, the final coherence order of –1 corresponds to the detected signal. The total crushing distance in k-space for each pathway is compared for the two gradient schemes (blue and orange). The optimization weighting assigned to each pathway is represented by the background of the plot. As shown, the new optimized gradient scheme achieves a larger crushing moment for most unwanted CTPs than the other approach, and especially for likely CTPs, demonstrating its potential for more effectively minimizing OOV artifacts.

**Figure 4:**
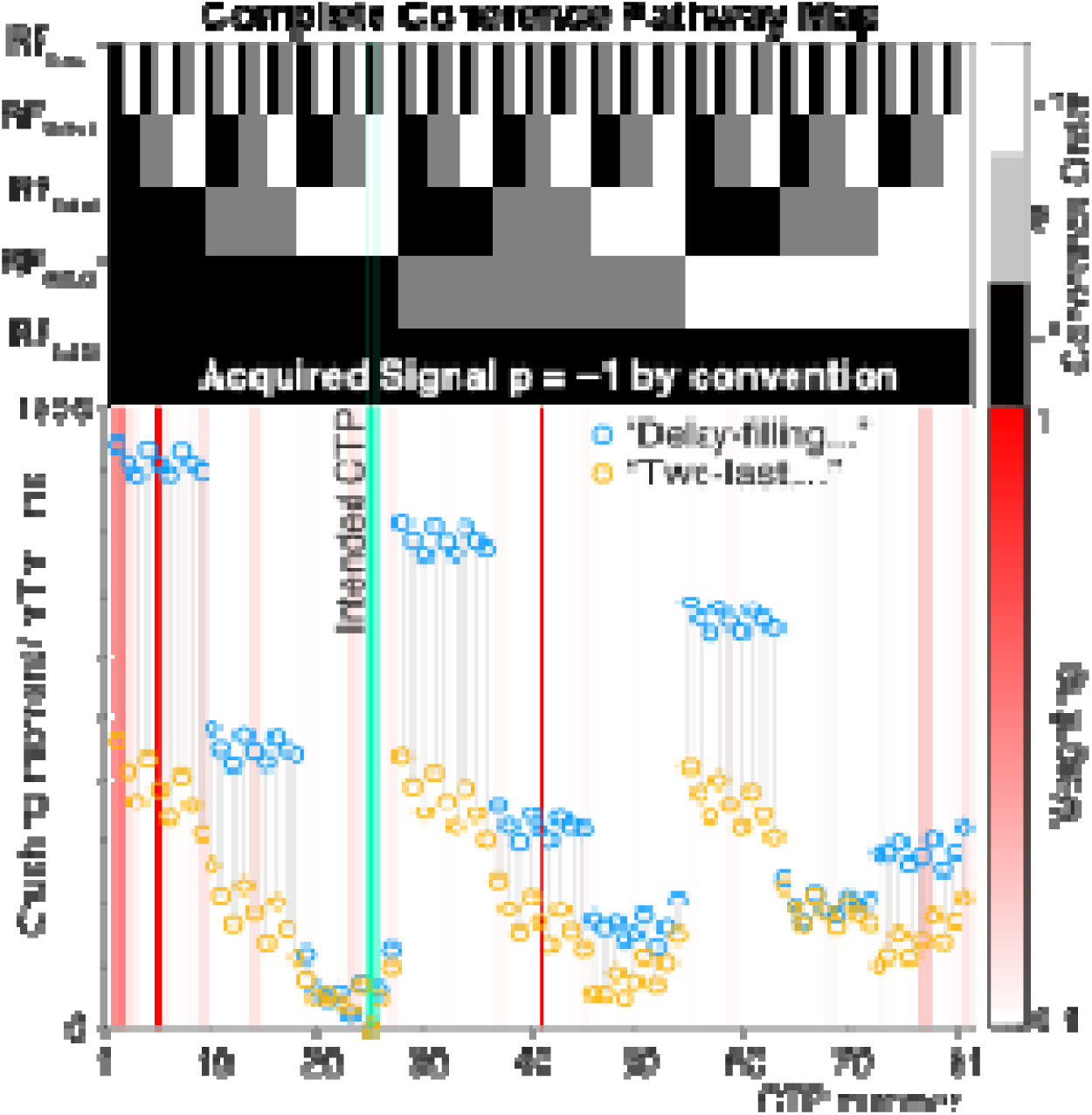
Comparison of total crushing moments of each coherence transfer pathways (CTP) for the “delay-filling optimized” and “two-last, increased-area” gradient schemes. The top section shows the complete CTP map, where each column represents one of the 81 possible pathways and each row shows the coherence order after an RF pulse. Coherence orders are color-coded: +1 in white, 0 in gray, and –1 in black. Note: Detectable signals are assigned a coherence order of p = −1 by convention. Total crusher moments (in mT·m⁻ ¹·ms) are plotted for each scheme, “delay-filling optimized” in blue and “two-last, increased areas” in orange. CTP #25 describes the intended signal pathway for this sequence. The excitation pulse alters the coherence order from 0 to −1. Subsequently, the first refocusing pulse changes the order from −1 to +1. The first editing pulse causes no change, leaving the order at +1. The second refocusing pulse then changes the order from +1 to −1, and again, the second editing pulse causes no change. A final coherence order of −1 is detectable. The background shading shows the weightings of each CTP. The “delay-filling optimized” scheme clearly increases crushing moments for most unwanted CTPs, suggesting it may more effectively suppress unwanted signal pathways.

### 3.2. Comparison of *in vivo* data

Representative spectral comparisons between the proposed “delay-filling optimized” and the “two-last, increased-area” scheme across three brain regions: PCC, thalamus, and mPFC, are shown in Figure 5. For each region, the SUM, GABA-edited, and GSH-edited difference spectra are shown alongside their respective model fits and residuals. In the thalamus (Figure 5A) and mPFC (Figure 5B), the “delay-filling optimized” scheme clearly demonstrates reduced OOV artifacts, as indicated by lower residual signals, particularly around 4.3 ppm. This improvement shows the effectiveness of the delay-filling optimization in minimizing OOV contamination, particularly in regions prone to susceptibility-related artifacts.

**Figure 5:**
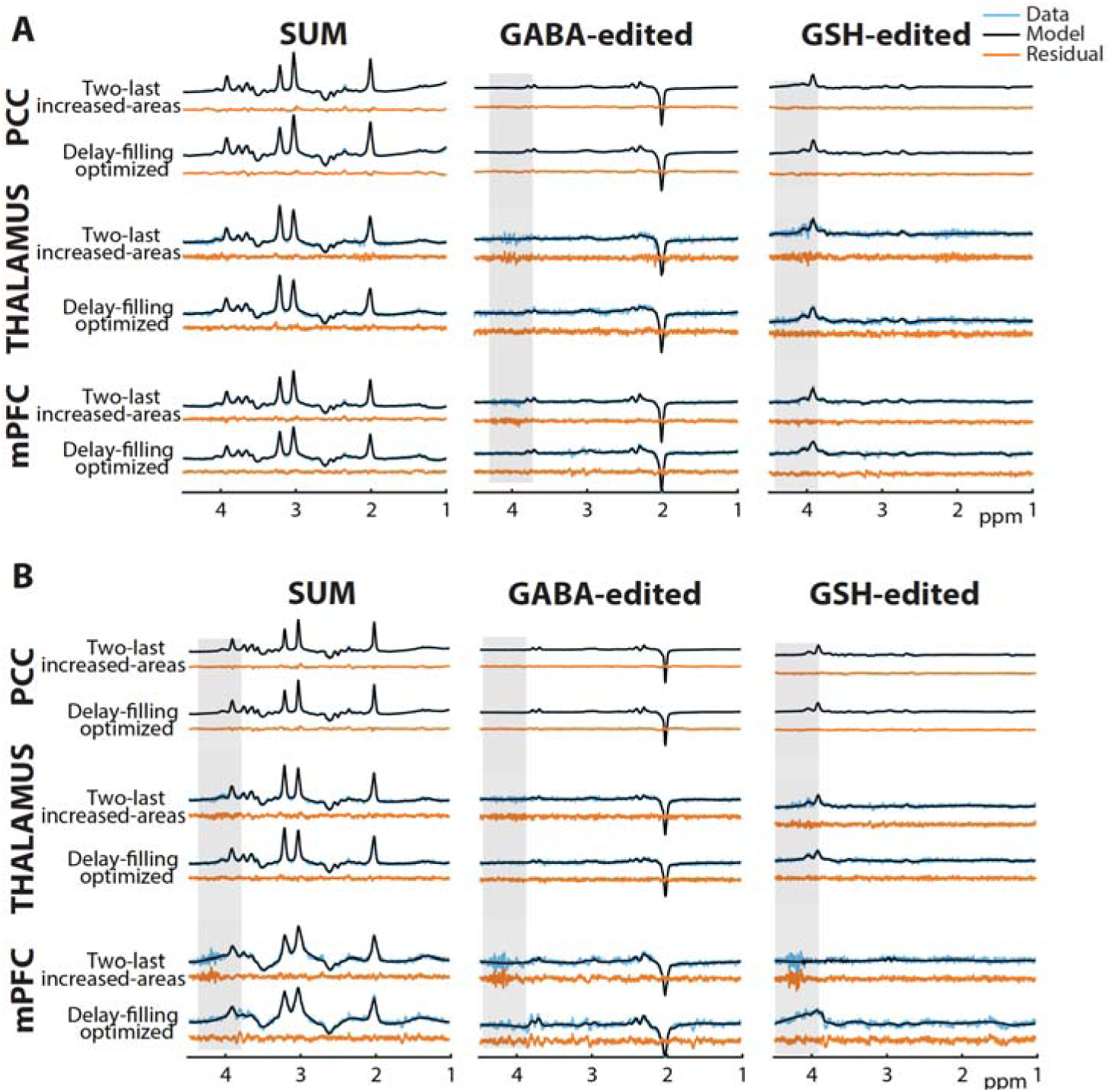
Spectra and model-residual comparisons between the proposed “delay-filling optimized” gradient scheme and the “two-last, increased-area” gradient scheme, using data from two representative volunteers across three brain regions: PCC, thalamus, and mPFC. Each panel shows the SUM spectrum, GABA-edited, and GSH-edited difference spectra (blue), their respective model fits (black), and residuals (orange). The delay-filling optimized gradient scheme shows reduced OOV artifacts in the thalamus (Figure 5A) and mPFC (Figure 5B) compared to the alternative scheme (highlighted in gray).

### 3.3. OOV artifact quantification

The relative amplitude of OOV residuals across subjects and brain regions is compared using box plots in Figure 6 and using a three-way ANOVA in Table 1. As shown in Table 1 there is a strong main effect of gradient scheme (*p* < 0.001), indicating that the proposed delay-filling optimized scheme significantly reduced artifacts compared to the other scheme. There was also a significant effect of subject (*p* < 0.001), showing expected variability in data quality based on individual anatomy. In contrast, voxel location did not have a significant overall impact (*p* = 0.248). A significant interaction was found between subject and voxel location (*p* < 0.001), and a three-way interaction among subject, location, and gradient scheme was also significant (*p* = 0.048), pointing to some variability in how the new scheme performed depending on both the individual and the voxel location. Overall, these results support the effectiveness of the optimized gradient scheme in reducing OOV-related artifacts.

**Figure 6:**
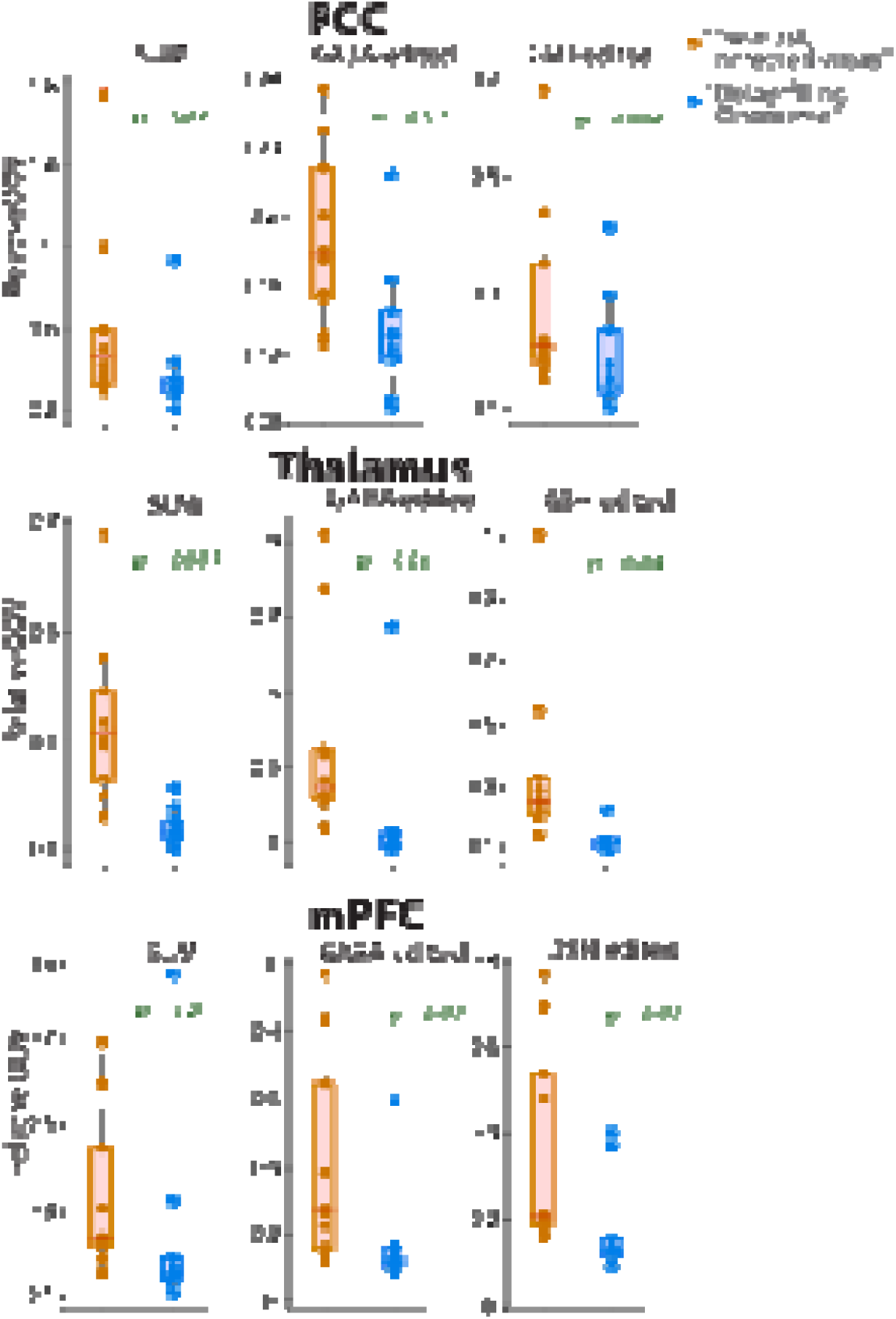
Comparison of relative OOV values (model residuals) across 10 datasets using the “two-last, increased-areas” scheme (orange) and the “delay-filling optimized” scheme (blue). Each data point (filled circle) represents the relative OOV values from an individual dataset. The p-values in the figure show the significance levels from a paired t-test of the two gradient schemes in each region and sub-experiments.

**Table 1:** Summary of three-way ANOVA results evaluating the effects of subject, voxel location, and gradient scheme (delay-filling optimized vs. two-last, increased-areas) on the relative OOV. A significant main effect of gradient scheme indicates that the optimized scheme substantially reduced OOV artifacts. Significant interactions suggest variability in the effect across subjects and regions (Sum Sq. – Sum of Squares, Mean Sq. – Mean Square, F – F-statistic, p-value – Probability Value).

| Analysis of Variance (ANOVA) |  |  |  |  |  |
| --- | --- | --- | --- | --- | --- |
| Source | Sum Sq. | d.f. | Mean Sq. | $F$ | $p$-value |
| Subject | 1.43 | 9 | 0.16 | 5.50 | < 0.001 |
| Voxel location | 0.08 | 2 | 0.04 | 1.41 | 0.24 |
| Gradient scheme | 1.23 | 1 | 1.23 | 42.38 | < 0.001 |
| Subject x Voxel location | 2.04 | 18 | 0.11 | 3.91 | < 0.001 |
| Subject x Gradient scheme | 0.51 | 9 | 0.07 | 1.94 | 0.05 |
| Voxel location x Gradient scheme | 0.05 | 2 | 0.03 | 0.93 | 0.39 |
| Subject x Voxel location x Gradient scheme | 0.89 | 18 | 0.05 | 1.70 | 0.04 |

### 3.4. Gradient scheme-induced diffusion weighting

The incidental diffusion weighting introduced by the delay-filling optimized gradient scheme resulted in a total-value of approximately 831.21 s/mm². Based on this - value and the known apparent diffusion coefficients (ADC), 1.6 × 10⁻□ mm²/s for tNAA, 1.41 × 10⁻□ mm²/s for tCho, 1.5 × 10⁻ mm²/s for Glu, 1.44 × 10⁻□ mm²/s for tCr, and 7.8 × 10^-^□ mm²/s for water (32), we estimated the corresponding signal attenuation () for each. The results showed modest signal reductions for metabolites in the range of 11–12%. Water exhibited a higher attenuation, with an of 0.6. The estimated values agree with the experimental data acquired as shown in Figure 7.

**Figure 7:**
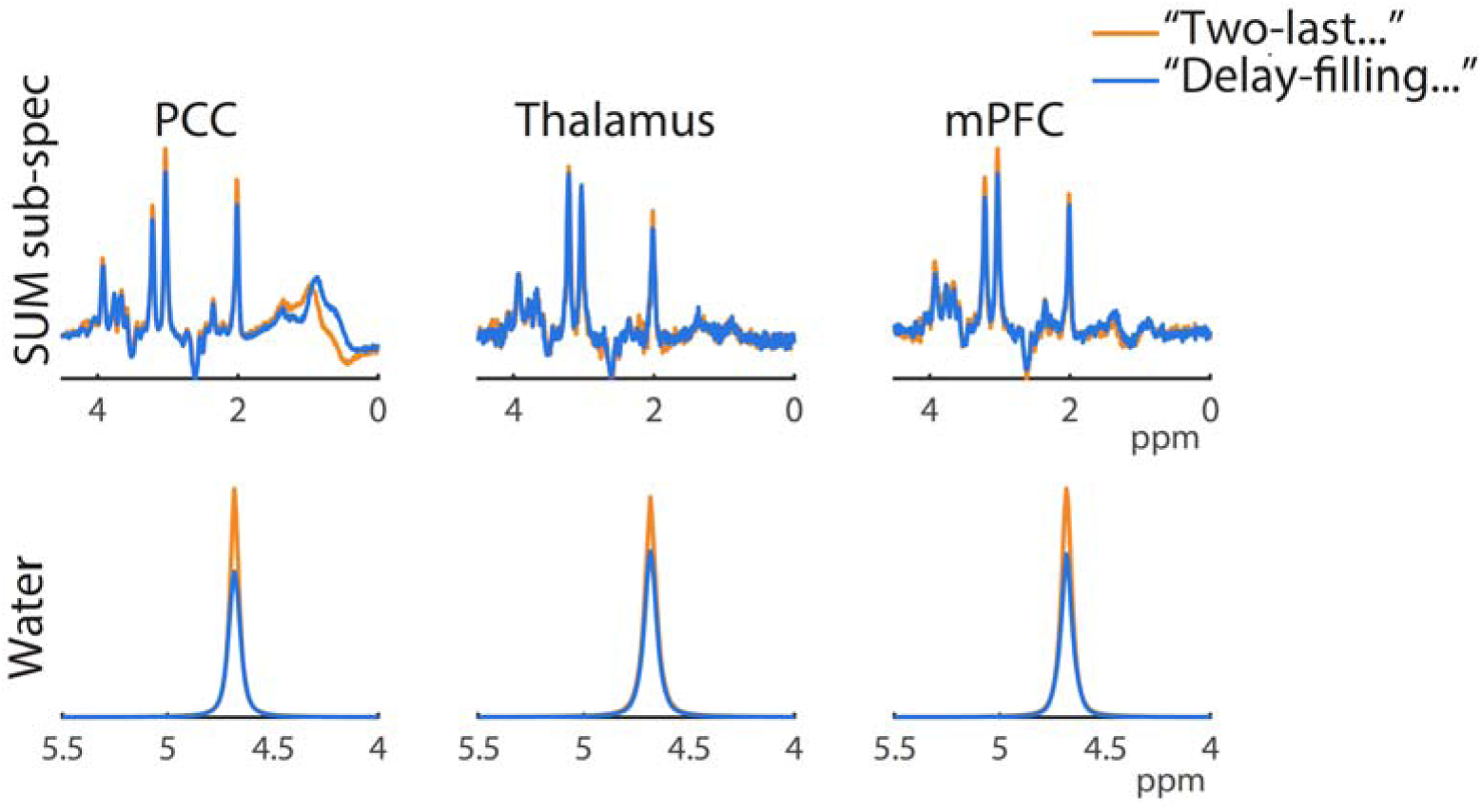
Comparisons of signal amplitudes for metabolite SUM sub-spectra and water acquired using the “two-last, increased-areas” and “delay-filling optimized” gradient schemes, illustrating signal losses due to diffusion.

## 4. Discussion

OOV artifacts are a long-standing challenge for single-voxel spectroscopy, obstructing metabolite signals and interfering with data processing algorithms. These unwanted signals are particularly pronounced in edited MRS due to the larger number of CTPs possible and the low amplitude of target signals, such as GABA and GSH. OOV signals originate from outside of the shimmed voxel of interest from pathways refocused by local field inhomogeneities. As each artifact arises from a specific coherence transfer pathway, it is critical to develop a tailored gradient scheme capable of suppressing coherence transfer pathways that are most likely to arise.

In this work, we presented a volume-based likelihood model for coherence transfer pathway weighting, integrated within a DOTCOPS-based gradient optimization framework for PRESS-localized edited MRS. Our approach aimed to prioritize suppression based on the estimated spectral contamination likelihood of each CTP, thereby enabling a more targeted and efficient use of gradient areas. A key feature of our model is the use of generalized, volume-based likelihoods to weight CTPs during optimization. The probability weights assigned to specific transitions (e.g., Δp = ±2 for a perfect refocusing pulse) were tied to the spatial and spectral regions defined by the selectivity profiles of the RF pulses. The values chosen (e.g., 10 for “out-of-slice”, 0.2 for “slice-edge”) are estimates intended to reflect the larger physical/spatial volume of tissue that does not experience an RF pulse (“out-of-slice/off-resonance”) compared to the precise transition bands of the pulses. This pragmatic approach prioritizes the suppression of signals originating from these large, outer volumes, which are a major source of OOV artifacts. While these probability estimates are simplified and do not represent precise calculations of a specific slice profile, they provide a tractable and effective mechanism to guide the optimizer towards gradient schemes that target the most probable sources of contamination, moving beyond a simple binary crushing paradigm. The cost function used for gradient scheme optimization has two terms: one that prioritizes suppression of the least-crushed CTP and one that prioritizes suppression on average. It is notable that the weightings are applied in this optimization in a different fashion than anticipated (but not used) in the DOTCOPS optimizer. In this implementation, important terms are emphasized in the mean by multiplied weights, and we make it more likely that a prioritized CTP would be considered by the minimum term by division of weights. In MEGA-edited PRESS sequences, the delays can have very different durations, and until now we have largely ignored this opportunity for increased CTP selectivity. The proposed model reflects a shift from the “one-size-fits-all” gradient approach (adapting MEGA-PRESS scheme only slightly from its parent PRESS) to a strategy that is more adaptive and efficient for OOV artifact reduction.

Using the proposed delay-filling and volume-based CTP likelihood weighting strategy, the genetic algorithm produced a scheme that maximized gradient areas across all three spatial axes with mixed polarity in the second axis. In terms of optimizer cost function, there are six variations on this scheme that have degenerate costs – each gradient axis independently can be inverted without changing the overall cost, as noted in the original DOTCOPS paper (20). While degenerate in terms of cost, these schemes are different in terms of the spatial origin of OOV signals that might arise (33). When evaluated in terms of total k-space crushing moment across CTPs, the “delay-filling optimized” scheme performed better than the “two-last, increased-area” approach, which was recently shown to perform better than our default.

These theoretical improvements were borne out in the *in vivo* data. Across three regions, the PCC, thalamus, and mPFC, the optimized gradient scheme showed reduced OOV artifacts, particularly around 4.3 ppm (as shown in Figure 5). Notably, the thalamus and mPFC, which are highly susceptible to *B*_0_ inhomogeneities and edge effects due to their proximity to air-tissue interfaces, showed more OOV signal and thus more opportunity for improved suppression, validating the robustness of the optimized scheme. The fact that voxel location did not exhibit a main effect (*p* = 0.248) suggests that the optimized scheme performs reliably across varied anatomical regions. Some inter-subject and voxel-scheme interaction effects were observed, which is not unexpected given known variability in head shape, positioning, and susceptibility distributions (34), The general trend favored the delay-filling optimized gradient scheme.

A key consideration in optimizing gradient schemes is the influence of diffusion effects, which can significantly impact metabolite quantification. Even within MRS sequences not primarily designed for diffusion measurements, some degree of diffusion sensitivity is inherently present due to gradients. Particularly for the maximalist scheme optimized here, which makes full use of gradient amplitude and duration limits, it is essential to quantify diffusion weighting through b-value calculations. In our study, the optimized crusher scheme introduced an effective b-value of approximately 831 s/mm², which resulted in an estimated 8–12% signal attenuation for metabolites. This level of diffusion weighting would require additional correction terms for absolute quantification. The observed diffusion attenuation was much stronger (∼40%) on the unsuppressed water reference signal, due to the more rapid diffusion of water. For quantitative MRS, it is common to acquire one water reference signal for eddy current correction (35) at long-TE, as here, and an additional reference at short-TE with less relaxation weighting. This second reference would be especially important with this degree of diffusion weighting. The Integrated Short-TE and Hadamard-edited Multi-Sequence (ISTHMUS) (36) automatically acquires long- and short TE water references, but for other sequences reference experiments are separately acquired.

To contextualize the performance of our proposed method, we selected the ‘two-last increased-areas’ scheme as the primary comparator, as it reflects our current best-performing in-house crusher scheme. This approach integrates our latest empirical understanding of problematic out-of-voxel pathways and incorporates practical adjustments to gradient timing and amplitude. By benchmarking against this baseline, we aimed to rigorously evaluate the specific contribution of the DOTCOPS optimization framework enhanced with volumetric weighting. A comparison against a vendor’s default scheme or an unweighted DOTCOPS version would certainly show improvement, but such a comparison would be less discerning, primarily indicating that the new method outperforms alternatives demonstrated to be sub-optimal elsewhere. Instead, by demonstrating superior artifact suppression relative to an already-refined scheme, we more convincingly establish the added value of our volume-informed, CTP-based optimization approach.

Although this study focused on PRESS-localized edited MRS, the optimized scheme could be extended to other sequences, such as STEAM or semi-LASER, which involve different coherence pathway dynamics and gradient requirements. In this work, we treated the underlying sequence timing as fixed and optimized the gradient areas within the available delays. MEGA-PRESS does have two degrees of freedom – the duration of the first and second slice-selective echo pulses, and the relative timing of slice-selective and editing pulses. These internal sequence delays could be varied within the optimizer in future work.

Additionally, the interaction between diffusion-induced signal attenuation and improved suppression could be modeled more explicitly in future optimization frameworks. Incorporating diffusion weighting into the algorithm, for instance by penalizing excessive signal attenuation during b-value estimation, could maintain effective OOV suppression with lower levels of diffusion weighting.

In the current implementation is that the optimized scheme requires a minimum TE of 80 ms. While this is suitable for GABA and GSH editing, it precludes its use for protocols requiring a shorter TE. However, the underlying optimization algorithm is general and can be applied to any set of timing constraints. Future work will focus on generating optimized gradient schemes for shorter TEs to broaden the applicability of the method. In addition, It is important to note that gradient performance parameters, such as the delay times and maximum amplitude, are vendor specific. While the optimization algorithm presented here can be used to generate platform-specific schemes, the experimental validation in this study was conducted on a Philips scanner. Future work will include implementation and validation on other major vendor platforms (e.g., Siemens, GE) to confirm the broader applicability of the proposed method.

## 5. Conclusion

In this study, we proposed and implemented an optimized gradient scheme for PRESS-localized edited MRS, using a volume-based likelihood model prioritizing CTPs based on spectral contamination likelihood. Using a genetic algorithm optimization and a delay-filling based approach, the resulting scheme demonstrated significant improvements in k-space crushing and suppression of unwanted pathways. *In vivo* validation across brain regions, including the PCC, thalamus, and mPFC, showed a reduction in OOV artifacts and improved spectral quality, even in regions prone to susceptibility challenges.

## Acknowledgements

This work was supported by NIH grants R01 EB016089, R01 EB023963, R01 EB032788, R01 EB035529, R21 EB033516, R00 AG02230, K99 AG080084, and P41 EB031771. GO is a paid consultant for Neurona Therapeutics Inc (unrelated to this work).

## Author Contributions

GLS contributed to Conceptualization, Methodology, Software, Validation, Formal Analysis, Investigation and Writing. ZS contributed to the Data Acquisition, Investigation, Resources and Data Analysis. SMM contributed to Data Acquisition, Writing, and Visualization. DS contributed to Data Acquisition, Writing, and Visualization. AG contributed to Data acquisition, Writing, and Visualization. YS contributed to Writing, and Visualization. VY contributed to the Investigation and Resources. CWD contributed to Writing, and Visualization. ATG contributed to Writing, and Visualization. HJZ contributed to Software, Formal Analysis, Writing, and Visualization. GO contributed to Software, Formal Analysis, and Supervision. RAEE contributed to Conceptualization, Methodology, Software, Validation, Formal Analysis, Investigation, Visualization, Writing, Supervision, Project Administration, and Funding Acquisition.

## Data availability statement

The data that support the findings of this study are available on request from the corresponding author. The data are not publicly available due to privacy or ethical restrictions.

## List of Abbreviations

^1^H MRS: Proton Magnetic Resonance Spectroscopy
ADC: Apparent Diffusion Coefficient
ANOVA: Analysis of Variance
Asc: Ascorbate
Asp: Aspartate
B_0_: Magnetic Field Strength (Static Field)
Cho: Choline-containing Compounds
CTP: Coherence Transfer Pathway
Cr: Creatine
D_av_: Average Diffusion Coefficient
DOTCOPS: Dephasing Optimization Through Coherence Order Pathway Selection
FIDs: Free Induction Decays
FH: Foot-Head (a spatial direction in MRI)
GABA: γ-Aminobutyric Acid
GA: Genetic Algorithm
Gln: Glutamine
Glu: Glutamate
GPC: Glycerophosphocholine
GSH: Glutathione
HERCULES: Hadamard Editing Resolves Chemicals Using Linear-combination Estimation of Spectra
HSVD: Hankel Singular Value Decomposition
mI: Myo-Inositol
ISTHMUS: Integrated Short-TE and Hadamard-edited multi-sequence
LCM: Linear-Combination Modeling
mPFC: Medial Prefrontal Cortex
MM: Macromolecule
MPRAGE: Magnetization-Prepared Rapid Gradient Echo
NAA: N-Acetylaspartate
NAAG: N-Acetylaspartylglutamate
OOV: Out-of-Voxel
PCC: Posterior Cingulate Cortex
PCh: Phosphocholine
PCr: Phosphocreatine
PE: Phosphoethanolamine
PRESS: Point-Resolved Spectroscopy
QM: Quality Metric
RF: Radiofrequency
sI: Scyllo-Inositol
STEAM: Stimulated Echo Acquisition Mode
Tau: Taurine
TE: Echo Time
TR: Repetition Time
VAPOR: Variable-Pulse Power and Optimized Relaxation
Δp: Change in Coherence Order

